# Effects of destruxin A on the trehalose content, trehalase activity and their gene expression of *Bemisia tabaci*

**DOI:** 10.1101/2023.06.28.546866

**Authors:** Can Zhang, Jianling Guo, Shaukat Ali, Bao-Li Qiu

## Abstract

*Bemisia tabaci* (Gennadius) (Hemiptera: Aleyrodidae) is a serious pest of agricultural crops in different regions of the world. Destruxin A (DA) is one of 39 derivatives of destruxins, which are secondary metabolites produced by entomopathogenic fungi. Due to the high insecticidal activity, DA have great application potential in biological control of *B. tabaci*. In order to understand the impact of destruxin A on the energy substances of the global pest *B. tabaci*, we measured the changes in energy substance content of the *B. tabaci* adults treated with DA at LC_10_ and LC_50_ concentrations after 4, 8, and 12 hours. The results revealed that DA caused varying degrees of decline in trehalose, soluble sugars, and glycogen content, indicating its ability to influence the energy substance content of whiteflies. After DA treatment, the content of trehalose in *B. tabaci* decreased significantly, while the activity of trehalase, a specific hydrolase for trehalose, increased significantly. Further study of the expression levels of two genes (*BtTre-1* and *BtTre-2*) encoding trehalases in *B. tabaci* after DA treatment showed that the soluble trehalase gene, *BtTre-1*, was induced in both low and high concentrations of DA, while the membrane-bound trehalase gene, *BtTre-2*, did not show significant changes in expression level.

## 1. Introduction

As a living organism, insects require a large amount of energy to complete their life activities, most of which are stored in their internal energy substances, including carbohydrates and fats. In the carbohydrates of insects, trehalose, glycogen, and glucose are the basic forms of energy storage. Studies have shown that insecticides can cause changes in the energy substances in insects. For example, after injecting deltamethrin into *Locusta migratoria*, trehalose significantly reduced after 24 hours (Alaoui et al., 1994). When injecting a high dose (LD_75_) of deltamethrin at 15 min and 6 h, glycogen storage in the fat bodies decreased by 100% (Alaoui et al., 1994). After treatment with DDT, the concentration of glycogen and trehalose in *Periplaneta americana* decreased rapidly in a short period of time (Granett et al., 1971). After being treated with triazophos, the trehalose content in the 5th instar nymphs of both long-winged and short-winged brown planthoppers decreased significantly (Ge et al., 2011). In addition to affecting carbohydrate content, the use of insecticides can also affect fat and protein. For example, carbosulfan can cause an increase in the accumulation of triglycerides and a decrease in the consumption rate of soluble proteins in *Liposcelis entomophila* and *L. bostrychophila* (Cheng et al., 2005). However, the impact of protein A disruption on the energy substances inside insects is currently unknown.

Trehalose exists in almost all tissues and organs in insects, especially in the hemolymph with the highest content. As the most important energy substance, trehalose plays a vital role in the energy metabolism of insects (Thompson, 2003). Trehalose has multiple physiological functions, mainly including: providing basic energy for the physiological activities of insects by decomposing to generate glucose; regulating the growth and development of insects (Shukla et al., 2015; Thompson and Redak, 2000); enhancing the anti-stress ability of insects under non-biological stress, such as protecting insect cells from damage caused by hypoxia (Chen et al., 2003); regulating the biosynthesis of chitin (Cohen, 1987). In addition, trehalose can also prevent the body fluids of insects from freezing and avoid frost damage (Thompson, 2003).

Trehalose is formed by the α,α-1,1 glycosidic linkage of two glucose units (α-d-glucopyranosyl-α-d-glucopyranoside)(Vinciguerra et al., 2022). Trehalase is the only enzyme that can specifically hydrolyze trehalose into glucose in the insect body. Therefore, trehalase is one of the most important enzymes in insect physiological processes, and changes in its activity will affect insect metabolism. Two types of aphid trehalase have been identified in insects, including water-soluble and membrane-bound enzymes(), which are encoded by *Tre-1* and *Tre-2*, respectively. Water-soluble trehalase mainly exists in the digestive and circulatory systems of insects, while membrane-bound trehalase mainly exists in muscles and plays an important role in the flight process of insects such as brown planthopper and locusts (Becker et al., 1996; Zhao et al., 2011). Currently, although much research has been done on disruptors, their effects on insect trehalase have not been reported.

In order to investigate the effects of destruxin A on the energy metabolism of Bemisia tabaci, this study examined the changes in carbohydrate content and trehalase activity in adult *B. tabaci* after treatment with low (LC_10_) and high (LC_50_) concentrations of destruxin A.

## 2. Materials and Methods

### 2.1. Insects and compounds

MEAM1 whiteflies were collected from cotton hosts and were reared at the Engineering Research Center of Biological Control of South China Agricultural University. The whitefly population was reared indoors on cotton plants (*Gossypium hirsutum*, Lu-Mian 32) in stainless steel rearing cages (60 cm × 60 cm × 60 cm) at a temperature of 26 ± 1□, relative humidity of 55 ± 10%, and a light cycle of 14 h light:10 h dark. Every 2 months, 20-30 adult whiteflies were randomly sampled from the population for purity testing to ensure a single population.

Destruxin A was extracted and purified from strain MZ16 of *Metarhizium anisopliae* isolated from MeiZhou, Guangdong referring to the method of Sree *et al*. (2015). DA sample had a purity of about 92%, including a small amount of destruxin A2, destruxin B, and some impurities. DA was diluted with DEPC water to a concentration of 10 mg/mL and assisted with a small amount of dimethyl sulfoxide (DMSO, Sigma) for solubility, and stored at -20□ for later use.

### 2.2. Destruxin A Treatments

The median lethal concentration (LC_50_) of DA was determined in a pilot experiment, in which *B. tabaci* adults were fed on six increasing doses of DA solution with 15% sucrose and 5% yeast powder. The dose that came closest to killing 50% of the adults (17.80 μg/mL) within 24 h was then selected for experiments.

After filtering and sterilizing with a 0.22 μm filter membrane, the solution was transferred to an artificial feeding device (transparent double-way tube) and covered with parafilm. 100 pairs of 3-day-old whitefly adults were transferred into the device from the other side and sealed with pierced parafilm. The feeding devices were wrapped in the lower half with black cloth and placed in a light incubator (26 ± 1°C, relative humidity 55 ± 10%, photoperiod L: D = 14 h: l0 h). After feeding for 4 h, 8 h, and 12 h, alive adults from each treatment were collected and stored at -80 °C. The control was fed by the feeding solution without DA, and each treatment was repeated 3 times.

### 2.3. Measurement of trehalose content

The 1-2 day-old whitefly adults, feed them with LC_50_ and LC_10_ concentrations of beauvericin respectively, and collect and process 100 *B. tabaci* (male and female ratio 1:1) after 4, 8, and 12 hours. Then, according to the ratio of 10 μL per pair of *B. tabaci*, add 500 μL of pre-cooled phosphate buffer (0.02 mol/L, pH 5.8), homogenize on ice with an electric grinder for 30 s, centrifuge at 4°C for 15 min, and take the supernatant as the measured homogenate. Trehalose content was estimated by the method designed by Steele et al. (1988) and Feng (1989).

### 2.4. Measurement of trehalase activity

The 3,5-dinitrosalicylic acid method was used to measure trehalase activity (Chrungu et al., 2017). The trehalase activity was calculated by measuring the glucose content generated by the decomposition of trehalose under the action of trehalase. The OD value was then measured at a wavelength of 550 nm using a microplate reader. The glucose generation was then calculated based on the standard curve, and the enzyme protein content was determined by the BCA protein concentration assay kit (Beyotime Biotechnology). Trehalase activity was determined as μmol glucose/mg protein/min.

### 2.5. Effects of destruxin A on soluble sugars and glycogen contents

Following the anthrone method of Halhoul and Kleinberg(1972), the soluble sugar and glycogen contents in *B. tabaci* were estimated. Adult *B. tabaci* of 1-2 days old were fed with LC_10_ and LC_50_ concentrations of beauvericin for 4 h, 8 h, and 12 h. After treatment, 100 *B. tabaci* (male and female in a 1:1 ratio) were collected and homogenized in 500 μL of pre-chilled phosphate buffer (0.02 mol/L, pH 5.8) with an electric grinder for 30 s on ice. The homogenate was then centrifuged at 10000 g at 4L for 10 min. The supernatant was transferred to a 2 mL centrifuge tube and mixed with 20% sodium sulfate solution (20 μL) and 1500 μL of chloroform-methanol mixture (chloroform:methanol=1:2) for 15 min at 10000 g at 4□ as the sample for determination. Measure the OD value at 630 nm wavelength using an enzyme-linked immunosorbent assay reader and use glucose as a standard.

### 2.6. Total RNA isolation and cDNA preparation

DA was diluted to LC_10_ and LC_50_ concentrations using ddH_2_O, and the 1-2 day-old whitefly adults were fed using a feeding device for 4, 8, and 12 hours. The control group was fed on liquid without DA. After the treatment, approximately 100 surviving adults were collected. Total RNA was extracted using Trizol reagent (Magen Biotechnology Co., Ltd, China), and the integrity, purity, and concentration of the RNA were verified by Agilent 2100 Bioanalyzer (Agilent, United States). cDNA synthesis was performed using the quantitative PCR reverse transcription kit, PrimeScript ™ RT reagent Kit (Perfect Real Time), and stored at -20°C. Each treatment was repeated three times. 100 *B. tabaci* adults were used for RNA extraction.

### 2.7. Quantitative Real-Time PCR (qRT-PCR)

Eighteen different genes were randomly selected from the detection results and verified by qRT-PCR of MEAM1 *B. tabaci*. The primers were shown in Table 1 and EF1α was selected as the internal reference gene. The cDNA (template) was obtained by using PrimeScript™ RT Reagent Kit (TaKaRa, Japan). The qRT-PCR reaction includes 10 μL SYBR qPCRMix (TOYOBO, Japan), 0.3 μM each primer, 7.5 μL sterilized water, and 2μL cDNA. The program was as follows: 95°C for 3 min, 40 cycles of 95 °C for 15 s, 60 °C for 30s, and 72 °C for 45s. The 2^-ΔΔCt^ technique was used to calculate relative expression levels using *BtTre-1* and *BtTre-2* primers and EF1α as an internal control (Wang et al., 2014). Three biological replicates were performed for each treatment, and three technical replicates were performed for each sample. All the experiments were performed with the Bio-Rad CFX-96 system.

### 2.8. Statistical analysis

For the changes in the activity of detoxifying enzymes in green peach aphids under different concentrations and treatment times, one-way analysis of variance (ANOVA) was performed using the statistical software SPSS (version 19.0) to compare the significance of the differences between each treatment. A significance level of *P* < 0.05 was taken, and after the variance analysis was tested for significant differences, Dunnett’s multiple comparisons were performed.

## 3. Results

### 3.1. Effect of protease A on trehalose content in green peach aphids

A standard curve of trehalose concentration and absorbance was plotted, and the regression equation was y = 0.1882x + 0.0037 with an R² value of 0.9971. Based on the standard curve, the trehalose content in green peach aphids treated with protease A was calculated, as shown in Figure 1. Significant decreases in trehalose content were observed after 4 hours of protease A treatment, and similar trends were observed at 8 and 12 hours. The lowest trehalose content was observed after 12 hours of treatment with an LC_50_ concentration, which was only 33.46% of the control. There was no significant difference in trehalose content between green peach aphids treated with LC_10_ and LC_50_ concentrations of protease A at the three time points (*P* > 0.05). These results suggest that protease A can significantly reduce trehalose content in green peach aphids.

**Figure 1.**
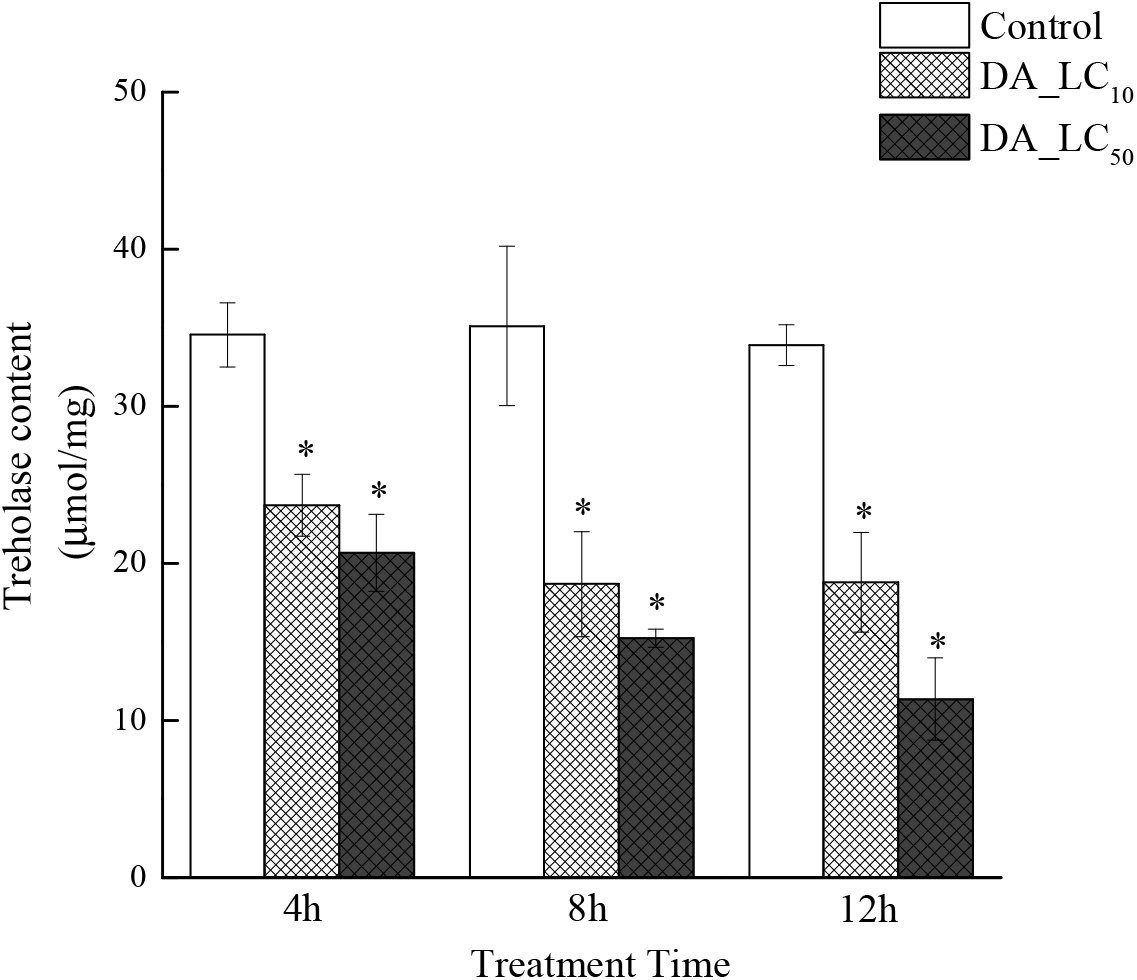
Changes of trehalose content in *Bemisia tabaci* after destruxin A treatment. (Values are the mean ± SE. * indicates significant differences from the control, *P* < 0.05)

### 3.2. Effects of protease A on the soluble sugar content of B. tabaci

A glucose standard curve was plotted with glucose concentration and absorbance as the x and y axes respectively, with a regression equation of y = 0.0373x + 0.0041 and R² = 0.9934.

The soluble sugar content of *B. tabaci* treated with protease A is shown in Figure 2. After 4 hours of treatment with low concentration protease A, there was no significant change in the soluble sugar content of *B. tabaci*, but high concentration protease A significantly reduced the soluble sugar content. After 8 and 12 hours of treatment with both low and high concentration protease A, the soluble sugar content of *B. tabaci* was significantly reduced, with no significant difference between the two concentrations. These results demonstrate a significant effect of protease A on the soluble sugar content of *B. tabaci*.

**Figure 2.**
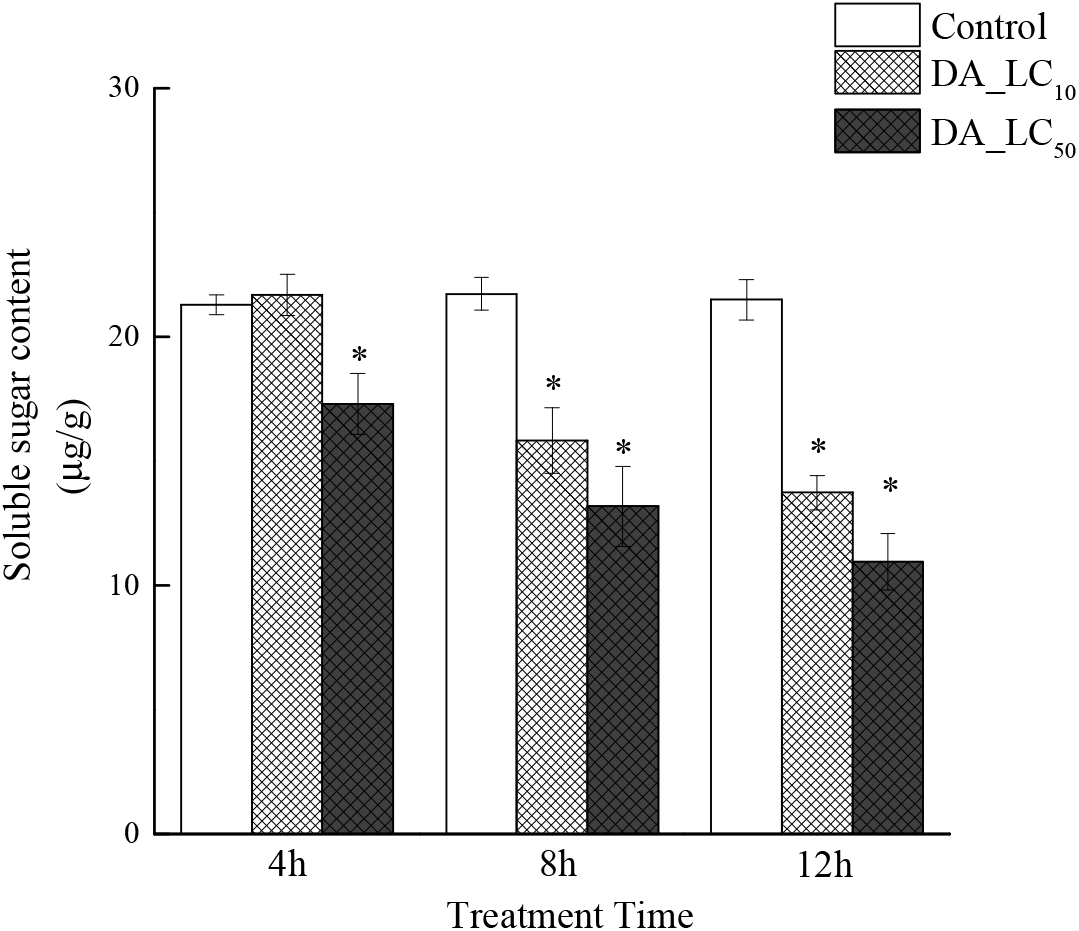
Soluble sugar content in *Bemisia tabaci* after destruxin A treatment. (Values are the mean ± SE. * indicates significant differences from the control, *P* < 0.05)

### 3.3. Effects of destruxin A on the Glycogen Content of the B. tabaci

As shown in Figure 3, DA treatment resulted in a significant decrease in glycogen content in the *B. tabaci* at both low and high concentrations, at various time points. The difference between the LC_10_ and LC_50_ concentrations was significant, with higher DA concentrations leading to a lower glycogen content in the *B. tabaci*. These results demonstrate that DA can significantly reduce the glycogen content of the *B. tabaci*, with a greater impact at higher concentrations.

**Figure 3.**
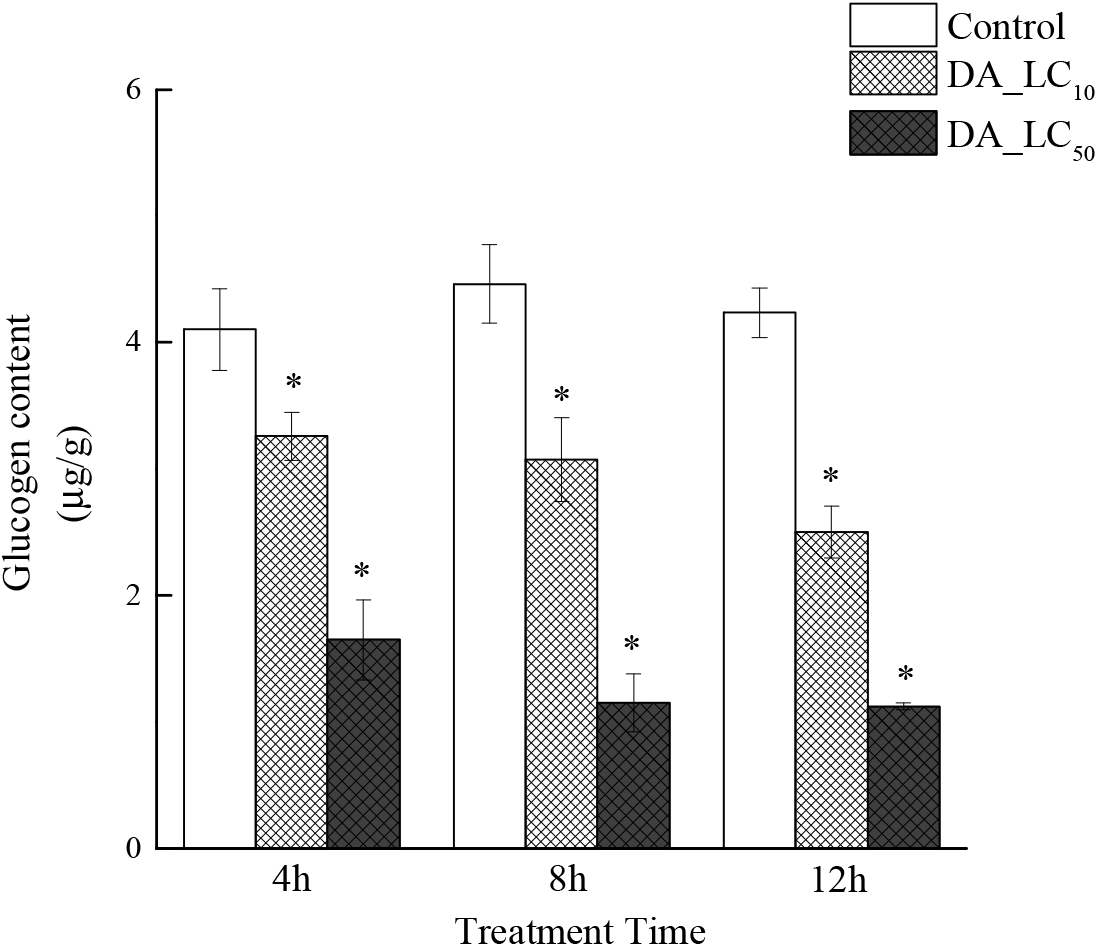
Changes of glucogen content in *Bemisia tabaci* after destruxin A treatment. (Values are the mean ± SE. * indicates significant differences from the control, *P* < 0.05)

**Figure 4.**
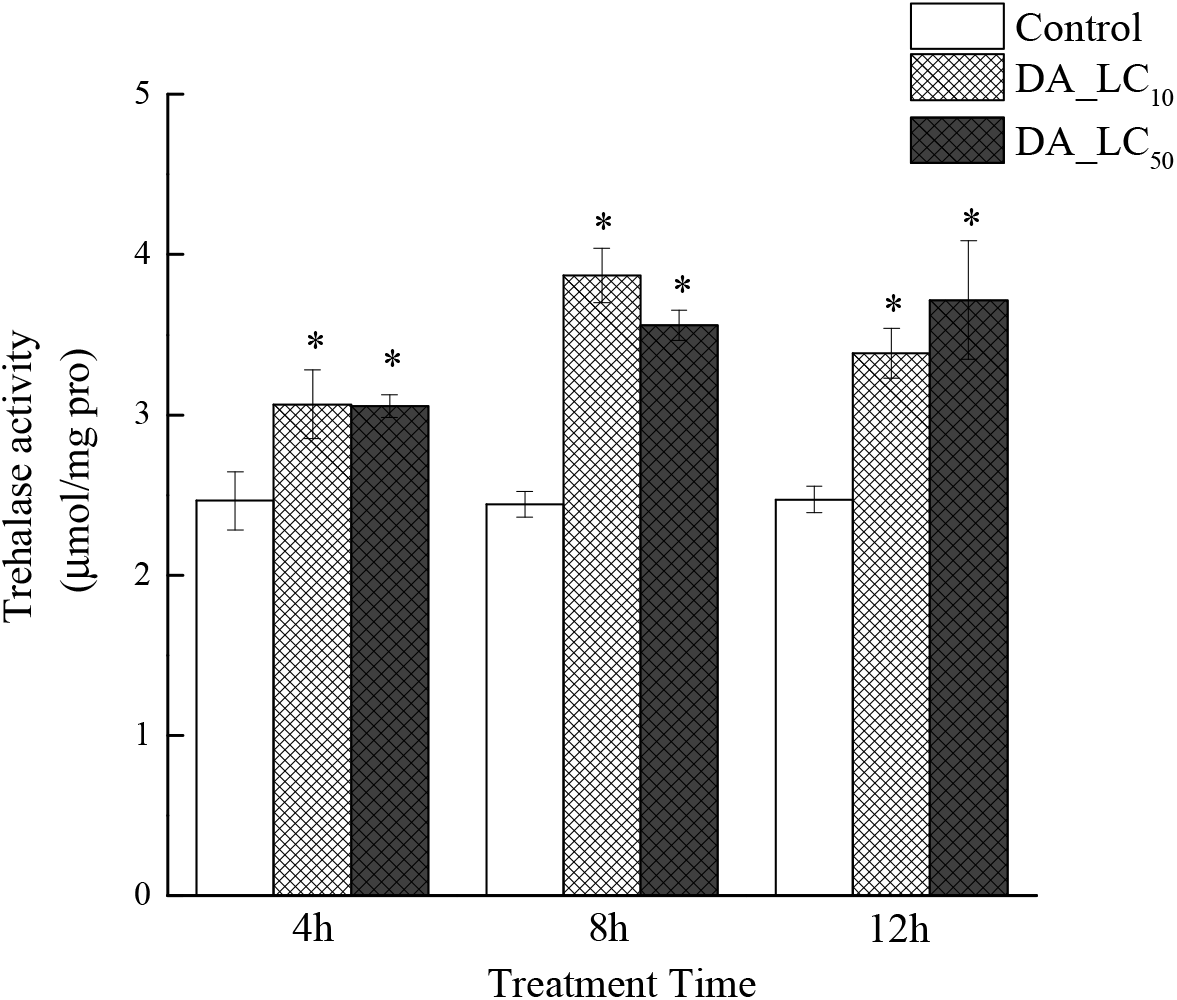
Changes of trehalase activity in *Bemisia tabaci* after destruxin A treatment. (Values are the mean ± SE. * indicates significant difference from the control, *P* < 0.05)

### 3.4. Effect of destruxin A on the expression level of trehalase gene

After treatment with sub-lethal concentration LC_10_ of DA, the expression level of the corresponding gene in the body of *B. tabaci* was shown in Figure 5. When *B. tabaci* were treated with LC_10_ concentration of DA, the expression level of *BtTre-1* in *B. tabaci* began to increase after 4 hours of treatment, significantly higher than the control level, reached the highest at 8 hours of treatment, and slightly decreased at 12 hours of treatment, but still significantly different from the control. However, the expression level of *BtTre-2* gene did not show significant changes at any time point.

**Figure 5.**
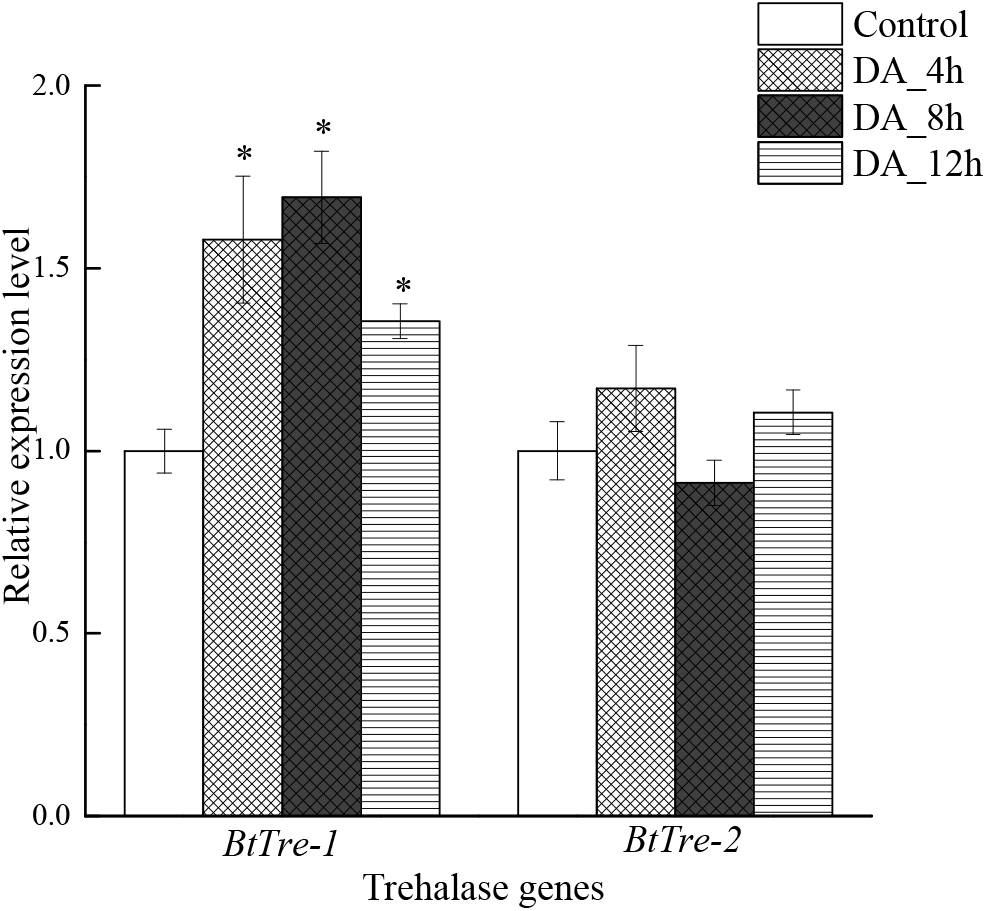
Relative expression of *BtTres* after treatment with destruxin A (LC_10_). (Values are the mean ± SE. * indicates significant differences from the control, *P* < 0.05)

After treatment with sub-lethal concentration LC_50_ of DA, the expression level of corresponding genes in the body of *B. tabaci* was shown in Figure 6. When *B. tabaci* were treated with LC_50_ concentration of DA, the expression level of *BtTre-1* in *B. tabaci* began to increase after 4 hours of treatment, maintained a high level at 8 and 12 hours of treatment, significantly higher than the control level. Similar to LC_10_ concentration treatment, the expression level of *BtTre-2* gene in *B. tabaci* did not show significant changes at any time point.

**Figure 6.**
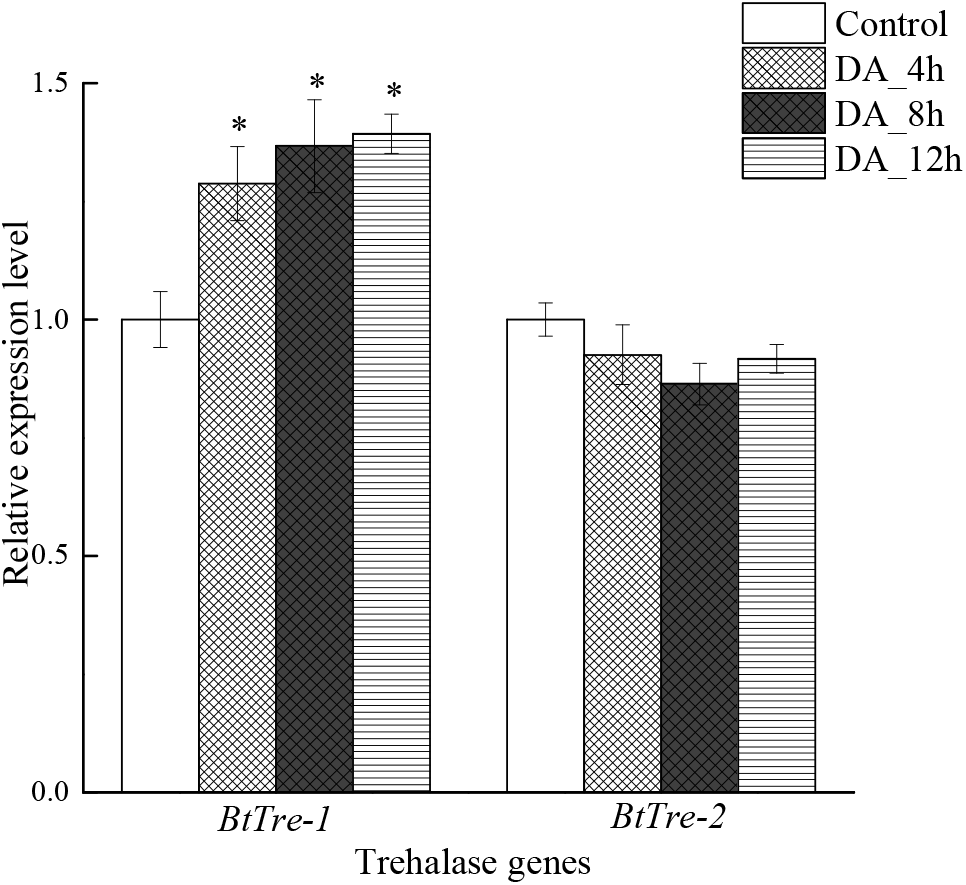
Relative expression of *BtTres* after treatment with destruxin A (LC_50_). (Values are the mean ± SE. * indicates significant difference from the control, *P* < 0.05)

## 4. Discussion

In order to understand the effect of destruxin A on the energy substances of the *B. tabaci*, we measured the impact of two concentrations of DA, including LC_10_ and LC_50_, on the levels of trehalose, soluble sugars, glycogen and other substances in the *B. tabaci*. The results showed that following treatment with abamectin, the levels of several carbohydrates in the *B. tabaci* decreased.

Trehalose is the most important carbohydrate substance in insects, commonly referred to as “blood sugar” and plays a crucial role in adaptation to non-biological stress (Shukla et al., 2015; Thompson, 2003). The concentration of trehalose affects insect growth and development. In this study, the concentration of trehalose in the *B. tabaci*’s body significantly decreased after treatment with DA, indicating that DA can affect the energy substance of the *B. tabaci*. In addition, studies have shown that trehalose helps protect cell membranes and proteins from damage or denaturation caused by various stress conditions (Elbein et al., 2003). The significant damage to the biological membrane system in the *B. tabaci* mentioned earlier after treatment with DA may also be related to the damage of trehalose.

Soluble sugars are important nutrient substances in insects, and insects can accumulate soluble sugars and other anti-stress substances under high temperature stress, thus increasing their own anti-stress ability (Ma et al., 2018). Therefore, when the level of soluble sugar in the *B. tabaci* significantly decreases, its own anti-stress ability is correspondingly reduced, accelerating the process of DA toxicity. In addition to soluble sugars, glycogen, as one of the energy storage substances in insects, also showed a decreasing trend in content after treatment with abamectin, indicating that DA may directly or indirectly affect the energy metabolism system of the *B. tabaci*. Previous studies have shown that DA has a strong repellency effect (Amiri et al., 1999; Thomsen et al., 2000). The results of this study showed that DA can significantly reduce the carbohydrate content in the body of *B. tabaci*, possibly because DA reduces the feeding ability of the *B. tabaci*, which in turn leads to a decrease in the carbohydrate content in its body, affecting the physiological and biochemical reaction processes of the *B. tabaci*. However, even after treatment with low concentrations of abamectin, the carbohydrate content in the *B. tabaci*’s body significantly decreased, indicating that the *B. tabaci* may have consumed a large amount of energy substances for metabolism, detoxification, and other related biochemical reactions.

In addition to trehalose content, destruxin A also had a significant impact on trehalase activity, showing that the trehalose content in the *B. tabaci*’s body significantly decreased, while trehalase activity significantly increased after DA treatment. Previous studies have shown that insecticides can affect changes in trehalase activity in insects. For example, the concentration of trehalose in male flying ants in East Asia significantly decreased after treatment with deltamethrin (Alaoui et al., 1994). Sub-lethal doses of imidacloprid, triazole phosphorus and bromocyanide pyrethroids can increase trehalase activity in brown planthopper, causing a decrease in trehalose content. This may be the physiological mechanism by which the flying ability of brown planthoppers is enhanced after insecticide treatment (Zhao et al., 2011). However, not all insecticides lead to increased trehalase activity in insects. For example, the plant-derived compound berberine has inhibitory effects on soluble trehalase in cotton bollworm intestinal tissues (Yu et al., 2016). In this study, the use of DA led to an increase in trehalase activity in the *B. tabaci*’s body, promoting the breakdown and metabolism of trehalose.

In order to study how destruxin A induces trehalase activity, we further measured the expression levels of two genes encoding trehalase in the *B. tabaci*’s body after DA treatment, and *BtTre-2*. The results showed that DA at both low and high concentrations could induce the expression of *BtTre-1*, but the expression level of *BtTre-2* did not significantly change. As *BtTre-1* is a soluble trehalase and *BtTre-2* is a membrane-bound trehalase, we speculate that DA only acts on soluble trehalase. Our results are consistent with Ge et al.’s results on brown planthoppers, where soluble trehalase activity increases after insecticide treatment, but membrane-bound trehalase activity does not change significantly, and the expression of the Tre-1 gene is up-regulated, while that of the *Tre-2* gene is not significantly different (Ge et al., 2011).

This study explored the consumption of carbohydrates in *B. tabaci* after treatment with DA and some of the reasons for this consumption. Although we cannot confirm whether the depletion of carbohydrates is the main cause of death in *B. tabaci*, the consumption of carbohydrates helps us understand the relevant biochemical reactions during the death process of the *B. tabaci* and may be directly or indirectly related to other factors characterizing DA poisoning.

